# Temporary knockdown of p53 during focal limb irradiation increases the development of sarcomas

**DOI:** 10.1101/2022.10.28.514234

**Authors:** Andrea R. Daniel, Chang-Lung Lee, Chang Su, Nerissa T. Williams, Zhiguo Li, Jianguo Huang, Omar Lopez, Lixia Luo, Yan Ma, Lorraine De Silva Campos, Sara R. Selitsky, Jennifer L. Modliszewski, Siyao Liu, Yvonne M. Mowery, Diana M. Cardona, David G. Kirsch

## Abstract

**Background:** Approximately half of all cancer patients receive radiotherapy and, as cancer survivorship rates increase with more effective therapies, the very low rate of radiation-associated sarcomas is rising. Radiation-associated sarcomas are life-threatening cancers, and radiation exposure is a primary risk factor for sarcoma development. During radiotherapy or other genotoxic cancer therapy for *p53* mutant cancers, pharmacological inhibition of p53 has been proposed to ameliorate acute injury of normal tissues. However, enhancing the survival of normal cells that sustain DNA damage by temporarily inhibiting p53 has the potential to increase the risk of cancer development. Here, we use *in vivo* shRNA technology to examine the consequences of temporarily reducing p53 expression on radiation-induced sarcoma development.

**Methods:** We utilized a mouse model of radiation-induced sarcoma where mice express a doxycycline (dox)-inducible p53 shRNA to temporarily and reversibly reduce p53 expression. Mice were placed on a dox diet 10 days prior to receiving 30 or 40 Gy hind limb irradiation in a single fraction and then returned to normal chow. Mice were examined weekly for sarcoma development and scored for radiation-induced normal tissue injuries. Radiation-induced sarcomas were harvested and subjected to RNA sequencing.

**Results:** Following single high-dose irradiation, 21% of temporary p53 knockdown animals developed a sarcoma in the radiation field compared to 2% of control animals. Mice with more severe acute injuries in the first 3 months after irradiation had a significantly increased risk of developing late persistent wounds in the soft tissue and bone. Chronic radiation-induced wounds were associated with sarcomagenesis. Examination of muscle stem cells by flow cytometry following hind limb irradiation indicated p53 knockdown preserves muscle stem cells in the irradiated limb, supporting the notion that temporary p53 knockdown at the time of irradiation reduces death of cells with DNA damage which may then persist to develop into a sarcoma. We performed RNA sequencing on 16 radiation-induced sarcomas compared to normal muscle controls. Gene set enrichment analysis revealed upregulation in the sarcomas of genes related to translation, epithelial mesenchymal transition (EMT), inflammation, and the cell cycle versus downregulation of genes related to myogenesis and tumor metabolism. Furthermore, genes with increased copy number such as *Met* and *Cdk4* were overexpressed in tumors.

**Conclusions:** Temporary reduction of p53 during high-dose irradiation increases late effects including tissue injuries and sarcoma development.

## Introduction

The p53 tumor suppressor protein is a critical component of the cellular DNA damage response machinery that drives cell death in acutely responding irradiated tissues (1). Following DNA damage, p53 initiates cell cycle arrest, apoptosis or senescence in a cell-type dependent manner (2, 3). Because p53-mediated cell death causes hematologic and other acute radiation and chemotherapy toxicities, temporarily blocking p53 during genotoxic therapies for patients with p53 mutant tumors has been proposed as a viable approach to prevent acute side effects of cancer therapy (1, 4). Preventing the death of cells injured by radiation is a strategy that has also been employed for the development of countermeasures to mitigate the acute radiation syndrome (ARS) following radiological disasters, theorizing that inhibiting programed cell death pathways will prevent the death of heavily damaged cells and preserve tissue integrity (5). While temporary inhibition of p53 or other cell death pathway components may spare sensitive normal tissues to radiation injury, it is also possible that by preventing the death of cells damaged by radiation late effects, including cancer development, may be exacerbated.

Approximately half of all cancer patients receive radiation therapy as part of their treatment (6). Radiation exposure, either from radiotherapy in the clinic, a radiation accident, or radiological warfare, is a primary risk factor for the development of sarcoma (7–10). Cancer survivorship rates are increasing as a result of improved cancer therapy, and as a corollary, rates of treatment-related malignancies are rising (11). Because children and young adults who survive cancer have many years to live after radiation therapy exposure, they are at an increased risk of developing radiation-induced sarcomas relative to older adults. Treatment-related secondary sarcomas are often aggressive and more challenging to treat than *de novo* tumors (8).

Sarcomas are life-threatening tumors that occur in adults, young adults, and comprise approximately 15% of all childhood tumors. Sarcomas are heterogeneous and aggressive malignancies that arise from the muscle, fat, bone, or other connective tissues. One of the most common childhood sarcomas, rhabdomyosarcoma, and one of the most common adult sarcomas, undifferentiated pleomorphic sarcoma (UPS), are tumors that can arise from muscle stem/progenitor cells or satellite cells (12).

Our previous work utilizing genetically engineered mice with *in vivo* shRNA targeting p53 demonstrated that temporarily reducing p53 expression in mice during fractionated low dose total-body irradiation reduced the risk of thymic lymphoma development (13). In this model, reduced p53-mediated apoptosis of bone marrow cells preserved cell competition in the thymic niche to prevent lymphoma-initiating cells harboring an oncogenic mutation in the thymus from developing into a lymphoma. Using the same genetically engineered mice with *in vivo* shRNA to p53, here we demonstrate that temporarily reducing p53 expression in mice during a single high dose fraction of radiation promotes sarcomagenesis through a cell autonomous mechanism.

## Materials and Methods

### Radiation-induced sarcoma mouse model

Radiation-induced sarcomas were induced as previously described (14). *CMV-rtTA; TRE-p53.1224* and *Actin-rtTA; TRE-p53.1224* mice were kindly provided by Scott Lowe (15). *CMV-rtTA; TRE-p53.1224* and *Actin-rtTA; TRE-p53.1224* mice express a doxycycline (dox)-inducible shRNA against p53 and their littermates harboring only a single allele (either *rtTA* or *TRE-p53.1224)* were used as controls (13). Mice were on a C3H and C57BL/6J mixed genetic background. Six to 24-week-old mice were placed on a dox diet for ten days, and then left hind limb of the mice was irradiated with a single fraction of 30 Gy or 40 Gy. Hind limb irradiation was performed using the X-RAD 225Cx small animal image-guided irradiator (Precision X-Ray). The irradiation field included the whole left hind limb and was defined using fluoroscopy with 40 kVp, 2.5 mA X-rays using a 2 mm Al filter (Figure S1). Irradiations were performed using parallel-opposed anterior and posterior fields with an average dose rate of 300 cGy/min prescribed to midplane with 225 kVp, 13 mA X-rays using a 0.3 mm Cu filter. Following irradiation, animals were immediately returned to normal chow.

After irradiation, mice were examined weekly for sarcomas. Upon detection, tumors were harvested with half stored in RNAlater (ThermoFisher Scientific) for subsequent RNA isolation and half formalin-fixed for histological analysis. Normal muscle samples were collected from contralateral (unirradiated) hind limbs from mice that did or did not develop sarcomas (Table S1).

### Satellite cell isolation and flow cytometry

Muscle satellite cells were isolated from *Pax7-nGFP (16, 17); Actin-rtTA; TRE-p53.1224* mice or littermate controls with *Pax7-nGFP* and only a single allele (either *Actin-rtTA* or *TRE-p53.1224).* Mice were fed dox diet for 10 days, irradiated with a single fraction of 30 Gy to one hind limb, and returned to normal chow. After 48 hours, mice were sacrificed and the muscles from the irradiated and unirradiated hind limbs were collected. Muscle satellite cells were isolated using a published protocol (18). Two million muscle cells were stained with Propidium Iodide and subjected to flow cytometric analysis to determine the percentage of live GFP+ muscle satellite cells in the irradiated and unirradiated limbs.

### Immunohistochemistry

Formalin fixed (10% neutral buffered formalin) paraffin embedded tumor tissues were sectioned (5 microns thick) and stained with hematoxylin and eosin. Immunohistochemistry with antibodies to S100 (Dako, GA504), Myod1 (Dako, M3512), Myogenin (Dako, IR06761-2), Desmin (Dako, M0760), Cytokeratin (Abcam, ab9377), CD31(Abcam, ab28364), SMA (Abcam, ab5694), and CD45 (BD Biosciences, 553076) were used to characterize tumor cell lineage and diagnosis. Sections from radiation-induced injured limbs were subjected to trichrome staining (Abcam, ab150686). Histology slides were reviewed by DC, an expert sarcoma pathologist, while masked to mouse genotype and treatment.

### Injury scoring

The irradiated limbs of mice were examined weekly and radiation-induced injuries were scored based on a system adapted from the Douglas-Fowler Skin Reaction Scoring system (19) as detailed in Table S2.

### RNA isolation and sequencing

Total RNA was isolated from tumors and normal muscle using Direct-zol RNA Miniprep Kit (Zymo Research). RNA samples were sequenced at the Duke Center for Genomic and Computational Biology Shared Resource. Total RNA was sequenced using paired end 150 base pair reads. 100 million reads were sequenced per sample in triplicate on a NovSeq instrument. RNA sequencing data upload to SRA (Sequence Read Archive) is in progress.

RNA-seq reads were aligned to mm10 mouse genome reference using STAR v.2.7.6a (20). Transcripts were quantified using Salmon v1.4.0 (21). Docker image used for alignment found on Dockerub: unclineberger/rna-seq-quant:2.3. Counts were normalized using DESeq2 (22). After normalization and transformation, counts were utilized in Principal Components Analysis for the purpose of identifying outliers. Genes were included in downstream analyses if they had at least 10 reads in any one sample. Differential expression analyses were performed with DESeq2. The Benjamini-Hochberg method was utilized to adjust p-values for multiple comparisons. Gene set enrichment analysis was performed with the mouse Hallmark (“mh”) and curated pathways (“m2”) gene sets obtained from Molecular Signatures Database (MSigDB) utilizing the fgsea package. Immune module gene sets from Charoentong et al. 2017 and Bindea et al. 2013 (23, 24) were converted to mouse gene symbols by sentence case conversion. The immune infiltration scores were computed by calculating median values for all genes in the set after scaling the normalized counts.

### Generation of p53/RB mutant tumors

The pX334-dual sgRNAs backbone vector was constructed by deletion of the Cre gene in the pX333-Cre vector (25) using two NcoI sites by standard cloning methods. For cloning pX334-sgTrp53, the pX334-dual sgRNAs vector was digested with BsaI enzyme and ligated to annealed sgRNA oligonucleotides targeting mouse Trp53 (Trp53 sgRNA: GTGTAATAGCTCCTGCATGG) (26). For cloning 4 individual pX334-sgTrp53-sgRb1 vectors, the pX334-sgTrp53 vector was digested with Bbs1 enzyme and ligated to annealed sgRNA oligonucleotides targeting 4 different loci of mouse Rb1 respectively (Rb1 sgRNA_1: AAATGATACGAGGATTATCG; Rb1 sgRNA_2: AGAGAAGTTTGCTAACGCTG; Rb1 sgRNA_3: TAAGTACGTTCAGAATCCAC; Rb1 sgRNA_4: GCAGTATGGTTACCCTGGAG) (27). Next, 50 ug of 4 equally mixed pX334-sgTrp53-sgRb1 plasmids were intramuscularly injected into at least six-week-old Rosa26^LoxP-Cas9-EGFP/LoxP-Cas9-EGFP^ mice with constitutive expression of Cas9 through *in vivo* electroporation as previously described (25). After injection, mice were examined weekly for sarcomas. Upon detection, tumors were harvested with half submerged in RNAlater (ThermoFisher Scientific) for subsequent RNA isolation.

## Results

### Temporary reduction in p53 expression during irradiation increases sarcomagenesis

We and others previously demonstrated that a single fraction of high dose irradiation (10-70 Gy) to the hind limb of wild type mice can lead to radiation-induced sarcomas in a small percentage of mice (14, 28). Furthermore, sarcomas can develop following low dose irradiation (1-4 Gy in single fraction) in mice deficient in p53 (29, 30). Notably, the p53 pathway is inactivated at a high frequency in human radiation-associated tumors (31), which suggests that loss of p53 function may play an important role in initiation and/or maintenance of radiation-associated tumors. To investigate if p53 function during irradiation plays an important role in initiation of radiation-induced sarcomas, we utilized a doxycycline (dox) inducible p53 shRNA mouse model to determine whether temporary knockdown of p53 during irradiation promotes sarcomagenesis (13, 14, 32). *CMV-rtTA; TRE-p53.1224* and *Actin-rtTA; TRE-p53.1224* mice or their littermate controls expressing either only the *rtTA* or *TRE-p53.1224* allele were fed a dox diet for 10 days to knockdown p53 expression and the hind limb was irradiated with a single fraction of 30 or 40 Gy. The mice were immediately returned to normal chow to restore p53 levels within 7 days (13) and followed for sarcoma formation for the entire natural course of their lives (schematic in Figure 1A). Of the 40 mice with p53 temporarily knocked down (p53KD) during 30 Gy irradiation, 8 developed sarcomas in the radiation field (20%), while no tumors were detected in the 48 control mice that received 30 Gy irradiation (Figure S2A). Similarly, 7 of the 32 (22%) p53KD animals that received 40 Gy to the hind limb developed in-field sarcomas compared to 2 of 49 (4%) control mice (Figure S2C). In total, 15 of the 73 (20.5%) irradiated p53KD animals and 2 of the 98 (2%) control animals developed radiation-induced sarcomas (Figure 1B). Notably, despite the increased development of sarcomas in the p53KD group, there was no significant difference in the overall survival between the p53KD and control groups at either radiation dose (Figure S2B and D). Additional cohorts of mice that did not receive dox were irradiated with 30 Gy to the hind limb to control for potentially aberrant expression of p53 shRNA independent of dox regulation. No significant difference was observed in sarcoma development within the radiation field between the mice harboring the two-allele system (*rtTA* and *TRE-p53.1224* alleles) for inducible p53 knockdown and the control mice with only one allele (either the *rtTA* or *TRE-p53.1224* allele) (Figure S2E). Furthermore, cohorts of mice harboring the two-allele or one allele system without irradiation were followed for their lifetime and no limb sarcomas were detected (Figure S2F).

**Figure 1.**
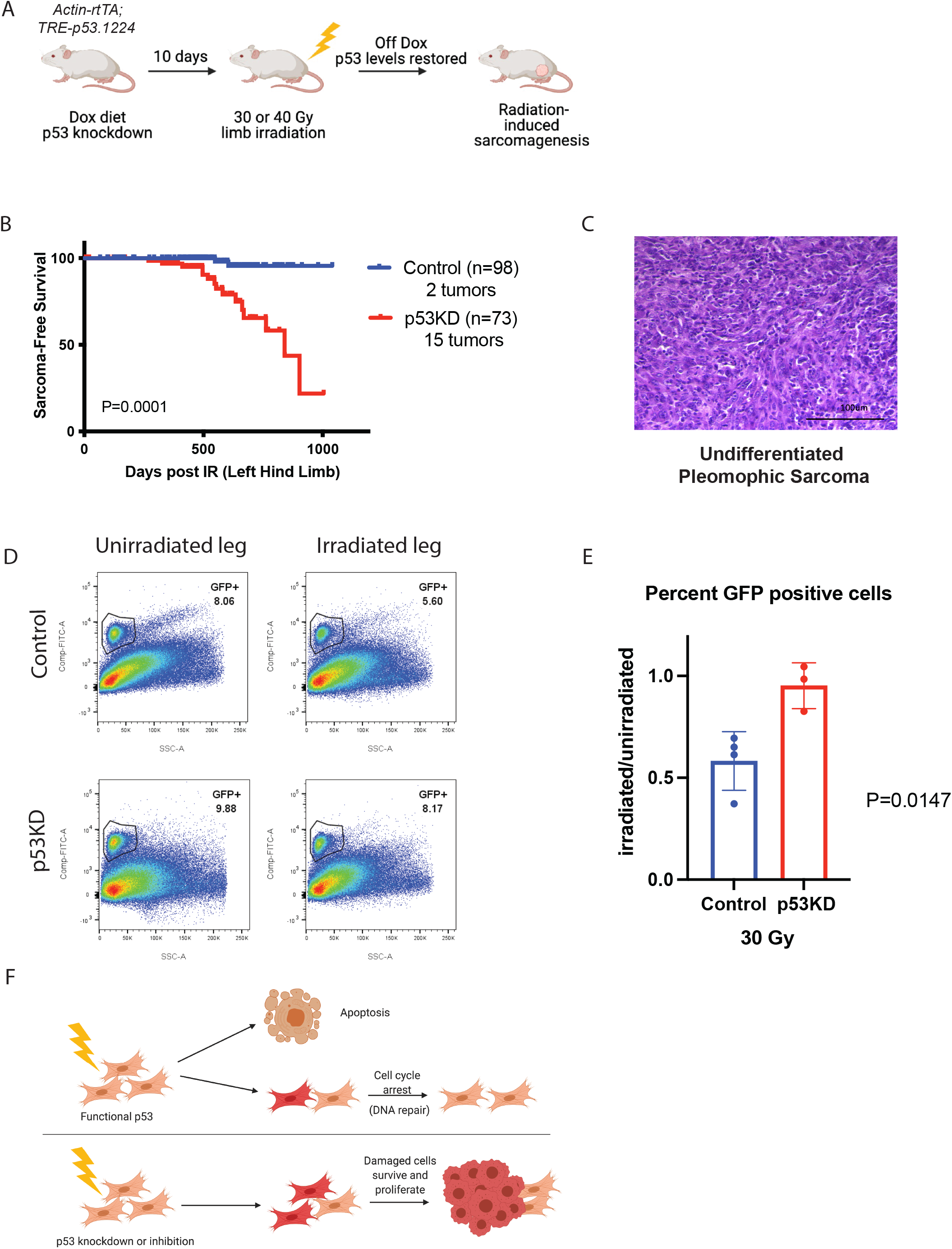
Temporary reduction in p53 expression during irradiation increases sarcomagenesis. (A) Schematic showing mice fed dox diet for 10 days to drive expression of shRNA to p53 to knockdown p53 (p53KD), irradiated with 30 or 40 Gy to the hind limb, returned to normal chow, and followed for sarcoma development and normal tissue injury in the radiation field. (B) Kaplan-Meier curves show radiation-induced sarcoma-free survival of control and p53KD mice irradiated with 30 or 40 Gy to the hind limb. P–value is from a log-rank test. (C) Representative image of a hematoxylin and eosin (H&E) stained radiation-induced undifferentiated pleomorphic sarcoma (UPS). (D) Representative flow cytometry dot plots of GFP+ muscle satellite cells isolated from unirradiated (left) and irradiated (right) limbs from control (top) or p53KD (bottom) mice. (D) The ratio of the percent of GFP+ cells in the irradiated limb over the unirradiated limb is graphed (±SEM). Each dot represents one mouse. P-value is from a T-Test. (F) Schematic showing that cells with functional p53 undergo apoptosis or cell cycle arrest and repair after irradiation, while damaged cells with impaired p53 function are protected from cell death and persist.

The radiation-induced sarcomas developed in mice between 269 and 903 days after irradiation with a median latency of 553 days (Table S1). Review of the H&E stained tumor sections and corresponding immunohistochemical analysis revealed that 13 of the 17 sarcomas were classified as undifferentiated pleomorphic sarcomas (UPS) (Figure 1C), 2 were designated as UPS with myofibroblastic differentiation, 1 as a leiomyosarcoma, and 1 as an angiosarcoma.

### Temporary knockdown of p53 during irradiation preserves muscle satellite cells

In our prior work we performed whole exome sequencing (WES) on a cohort of radiation-induced sarcomas from this mouse model and determined that while these sarcomas exhibited a relatively low nonsynonymous somatic mutational burden compared to carcinogen-induced sarcomas, they exhibited a distinct genetic signature characterized by C-to-T mutations that is indicative of oxidative damage (14). The radiation-induced mouse sarcomas also displayed a high indel to substitution ratio and a frequent gene copy number variations (14) which is consistent with the types of genetic damage observed in human radiation-associated sarcomas (33, 34). These data led us to hypothesize that the increased rate of sarcomagenesis in the irradiated p53KD animals, compared to controls with wild type p53 levels, may be due to protection from radiation-induced p53-mediated cell death of tumor-initiating cells. Cells expressing wild type levels of p53 that exhibit extensive radiation damage may undergo p53-mediated apoptosis (35). When expression of p53 is temporarily reduced or inhibited during irradiation, the damaged cells may survive and initiate sarcomas. Because many of the radiation-induced tumors in our model are UPS tumors and because we and others have previously used Pax7-CreER mice to show that mouse sarcomas that mimic human UPS can arise from Pax7+ myogenic progenitor cells (12, 36), we examined the fate of satellite cells in mice with temporary p53 KD during irradiation. Satellite cells express the muscle lineage marker Pax7. To label muscle satellite cells in our model, we crossed the p53KD mice to Pax7-nGFP mice (16). Mice expressing nGFP in Pax7+ cells were placed on dox diet for 10 days to knockdown p53, irradiated with 30 Gy to one hind limb, and then returned to normal chow. After 48 hours, when satellite cells are ablated after high dose irradiation (37), muscles from the irradiated and the unirradiated hind limbs of each mouse were collected and muscle stem cells were isolated. Flow cytometry for GFP+ cells was performed to determine the relative percentage of live satellite cells remaining in the muscle 48 hours after irradiation (Figure 1D). The percent of GFP+ cells remaining in the irradiated leg compared to the unirradiated leg of each mouse was determined (Figure 1E). In the p53KD mice, there was minimal loss of GFP+ muscle satellite cells after irradiation, while approximately 40% of GFP+ muscle satellite cells were lost in the control mice. The survival of muscle satellite cells in the p53KD animals after high dose irradiation supports a model whereby the persistence of damaged irradiated cells, that would have undergone p53-mediated apoptosis, may initiate oncogenic transformation and eventual sarcoma formation (Figure 1F, schematic).

### Temporary reduction in p53 expression during irradiation promote chronic injuries

The strategy of inhibiting p53 during genotoxic therapies to reduce acute side effects may exacerbate other late effects of radiation in addition to carcinogenesis. The consequence of p53-mediated signaling in response to radiation is cell- and tissue-type dependent. For example, loss of p53 may block apoptosis and/or promote mitotic death (13, 38). Therefore, the probability of developing late effects of hind-limb irradiation following p53 knockdown may vary with tissue type (ie. muscle, skin, vasculature, nerves, fat, and bone). Radiation-induced chronic wounds may occur due to an acute wound that fails to heal or may arise months to years after radiation exposure in tissue that initially appears to have recovered from or avoided acute toxicity (39). Late persistent wounds are characterized by inflammation, ulceration, fibrosis, and/or necrosis of soft tissue and bone. Damage to the vasculature of irradiated tissues may contribute to impaired wound healing due to a lack of neovascularization leading to insufficient perfusion (40). We previously showed that p53 is required in endothelial cells to prevent radiation-induced injury to the heart (41).

We observed acute and chronic injuries to the irradiated leg in the control and p53KD mice. Mice were evaluated weekly based on a previously published rubric for skin injury that we adapted to comprehensively assess radiation-induced normal tissue toxicity of the skin, bone, and muscle (19) (Table S2). Mice exhibiting signs of injury (skin breakdown and/or swelling) were given a score of 1 and scoring increased with the severity of the injury to a maximum score of 4 (loss of the foot). Acute injuries were defined as occurring within the first three months of irradiation, and late injuries were defined as arising after 3 months. Injuries were treated with topical antibiotics and supportive care, including wet food and hydration gel packs provided to injured mice. Late injuries presented as chronic wounds that slowly progressed. Therefore, with few minor exceptions, the last injury score was the highest score the mouse received.

Chronic tissue injuries were examined histologically by a sarcoma pathologist, masked to genotype and treatment, and the degree of fibrosis was assessed using H&E and trichome stained slides (Figure S3A-F). Tissue from animals scoring a 1-1.75 showed evidence of muscle and skin fibrosis, while those from animals with injury scores between 2 and 2.75 displayed muscle, skin, and neurovascular fibrosis. Half of the tissue injuries scoring 3-3.75 exhibited histological evidence of skin and muscle fibrosis with an increase incidence of neurovascular fibrosis and some bone remodeling. Animals with an injury score of 4 exhibited histological evidence of muscle, skin, and neurovascular fibrosis, and bone remodeling (Figure S3G-H). Evaluation of injuries in mice receiving 3+ scores from the p53KD and control groups reveal the groups to be similar with slightly more neurovascular fibrosis and slightly less skin fibrosis in the p53KD group (Figure S3I).

Examination of acute injuries scoring at least a 1 demonstrated a higher frequency of injuries in the p53KD group compared to control in the animals that received 40 Gy hind limb irradiation (Figure 2A), but no significant difference was observed in the animals that received 30 Gy (Figure 2B). In contrast, the development of chronic injuries was significantly greater in the p53KD animals that received 30 Gy compared to control animals for score levels 1-3, but not 4 (Figure 2C-F). For animals that received 40 Gy, a significant increase in chronic injuries in the p53KD group compared to control was only observed for the score level 2 (Figure S4A-D). The combined control groups exhibited a median final injury score of 1.375 compared to 2.375 from the p53KD animals, and this difference did not reach statistical significance (Figure S4E). Notably, in the control cohorts that did not receive dox prior to 30 Gy irradiation, no significant difference was observed between the genotypes at any score level (Figure S5).

**Figure 2.**
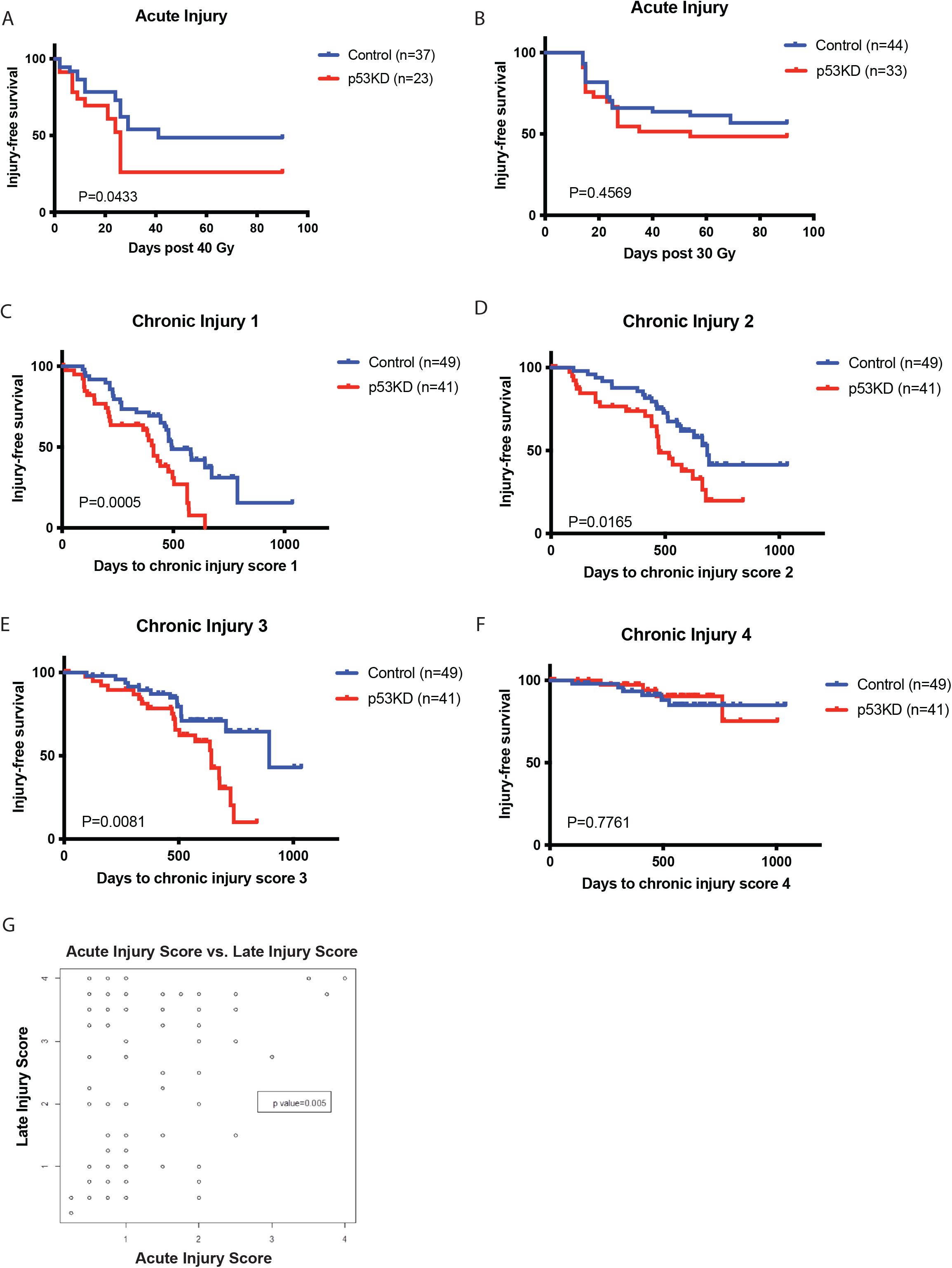
Temporary reduction of p53 during irradiation increases chronic injuries. (A) Kaplan-Meier curves show acute injury-free survival (score 1+) of control and p53KD mice irradiated with 40 Gy to the hind limb. P–value is from a log-rank test. (B) Kaplan-Meier curves show acute injury-free survival (score 1+) of control and p53KD mice irradiated with 30 Gy to the hind limb. P–value is from a log-rank test. (C-F) Kaplan-Meier curves show chronic injury-free survival from scores 1+ (C), 2+ (D), 3+ (E), or 4 (F) of control and p53KD mice irradiated with 30 Gy to the hind limb. P–value is from a log-rank test. (G) Correlation coefficient test compares acute injury scores and late (chronic) injury scores of the control and p53KD mice that received either 30 or 40 Gy to the hind limb. P-value was generated using a chi square test for the association between late score and acute score.

We next performed a correlation coefficient test comparing the acute and chronic injury scores from all the mice, which included the p53KD and control mice that received 30 or 40 Gy (Figure 2G). This analysis showed an association between mice with a high acute injury score and the development of a high chronic injury score. However, low acute injury score was not associated with low or high chronic injury score.

### Radiation-induced chronic injuries increase the risk of sarcomagenesis

Understanding the relationship between tissue damage and tumor promotion may improve secondary cancer detection and prevention strategies (42). We previously observed that muscle tissue injury promotes sarcoma development by stimulating muscle satellite cell activation (43, 44). In addition, chronic inflammation and wound healing can create a tumor promoting microenvironment with a permissive immune milieu (45–48) and epigenetic reprogramming to stimulate tumor outgrowth (49). We compared the final chronic injury scores of mice from the p53KD and control groups receiving 30 or 40 Gy that developed a radiation-induced sarcoma to those of mice that did not develop a sarcoma in the radiation field (Figure 3A). The sarcoma-free mice had a median chronic injury score of 1.5, while the median score of the sarcoma-bearing mice was significantly higher at 3.75. Of the 17 animals that developed radiation-induced sarcomas, 15 exhibited injury scores greater than 3. Animals in this group were observed to have a level 3+ injury for 29 to 455 days (median 132 days) prior to tumor detection. A Cox model analysis demonstrated that, independent of the covariants dose and genotype, radiation-induced chronic injuries are associated with sarcomagenesis (Figure 3B).

**Figure 3.**
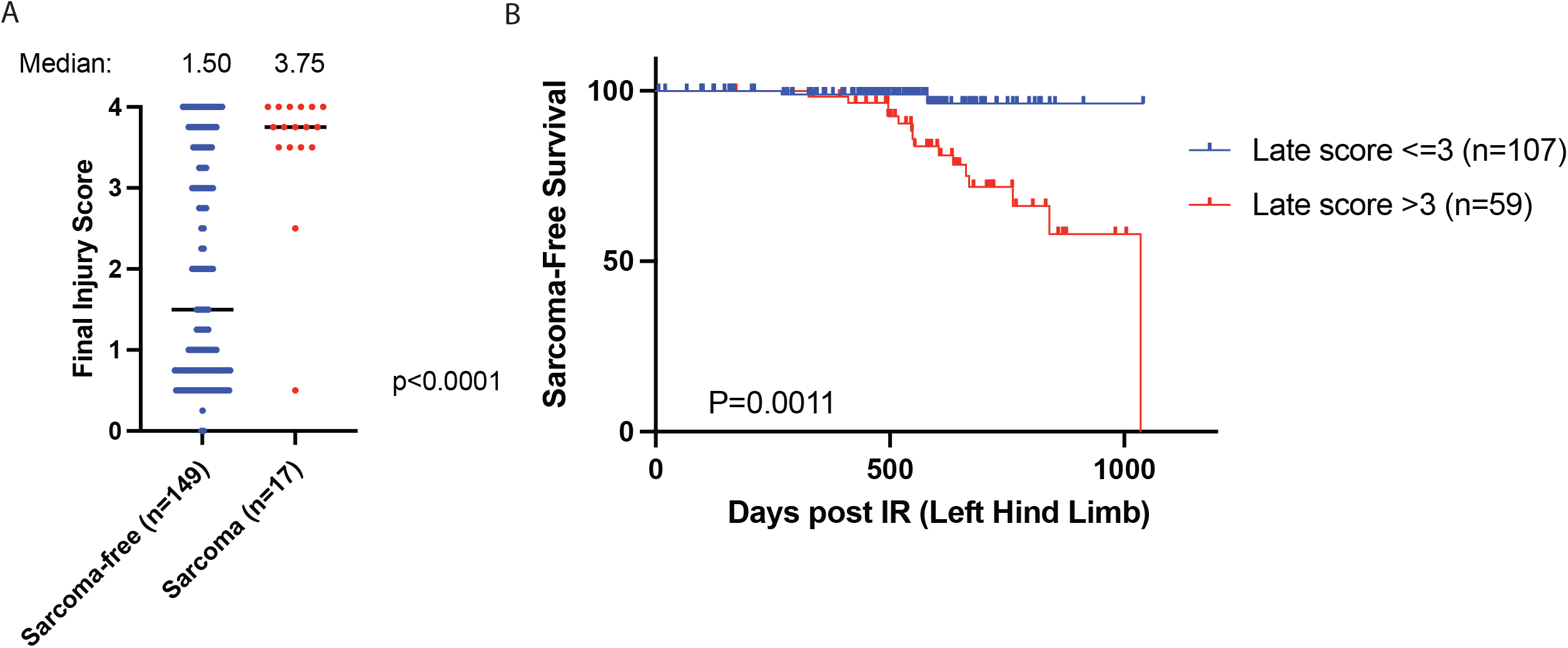
Radiation-induced chronic injuries increase the risk of sarcomagenesis. (A) The final injury scores of the control and p53KD mice that received 30 or 40 Gy are plotted. The final injury scores of mice that did not develop a radiation-induced sarcoma (blue) are compared to the scores of mice that did develop a radiation-induced sarcoma (red). P-value is from a T-Test. (B) Kaplan-Meier curves show radiation-induced sarcoma-free survival of the control and p53KD mice irradiated with 30 or 40 Gy to the hind limb. Mice with chronic injury scores equal to or greater than 3 are compared to mice with chronic injury scores less than 3. P–value is from a cox proportional hazard model.

### Radiation-induced sarcomas exhibit an increase in proliferative gene expression programs and a decrease in myogenic differentiation gene expression programs

In our prior analysis of WES data from radiation-induced sarcomas, no recurrent nonsynonymous somatic mutations in oncogenes or tumor suppressor genes were identified (14). To gain insight into the mechanisms of radiation sarcomagenesis, we analyzed RNAseq results from 16 radiation-induced sarcomas in our cohort and normal muscles from 7 age-matched littermate mice (117-840 days old, median age 343 days) (Table S1). As an additional control, we also performed RNAseq on two sarcomas initiated by loss of the *p53* and *Rb1* tumor suppressor genes. Principal component analysis revealed that the sarcoma samples clustered together, and the normal muscle samples clustered as a group (Figure S6A). Upon examination of the top 75 differentially expressed genes, the 14 UPS and one leiomyosarcoma clustered together (Figure 4A). Not surprisingly, based on transcriptomic analysis of human radiation-associated sarcomas (34), the angiosarcoma sample did not cluster with the other radiation-induced sarcomas. The radiation-induced sarcomas were characterized by a preponderance of gene downregulation events (n=1,051 Q<0.05) compared to upregulation of genes (n=657 Q<0.05, Figure 4B). The top downregulated gene was the muscle differentiation marker *MyoC* (50) (Figure S6B), and the top upregulated gene in the radiation-induced sarcomas was the cytokine *Wisp1* (Figure S6C), which can promote proliferation (51). Notably, the inflammatory cytokine *Il1f6* was among the most upregulated genes in the tumors compared to normal muscle (Figure S6D). Gene set enrichment analysis revealed that the genes upregulated in tumors were related to epithelial to mesenchymal (EMT) transition, inflammation, and cell cycle, while the downregulated pathways were related to myogenesis and metabolism (Figure 4C and S6E-F). The p53/RB sarcomas engineered to lack RB function serve as controls for activated Hallmark E2F target gene expression (Figure S6E). Analysis of immune infiltration in tumors indicated a general increase in immune response at the gene level, including significant enrichment of inflammatory cells including mast cells, macrophages, and some T lymphocytes (Figure S7). The tumors also exhibited enrichment of myeloid-derived suppressor cells and regulatory T cells indicating an immunosuppressive microenvironment (24).

**Figure 4.**
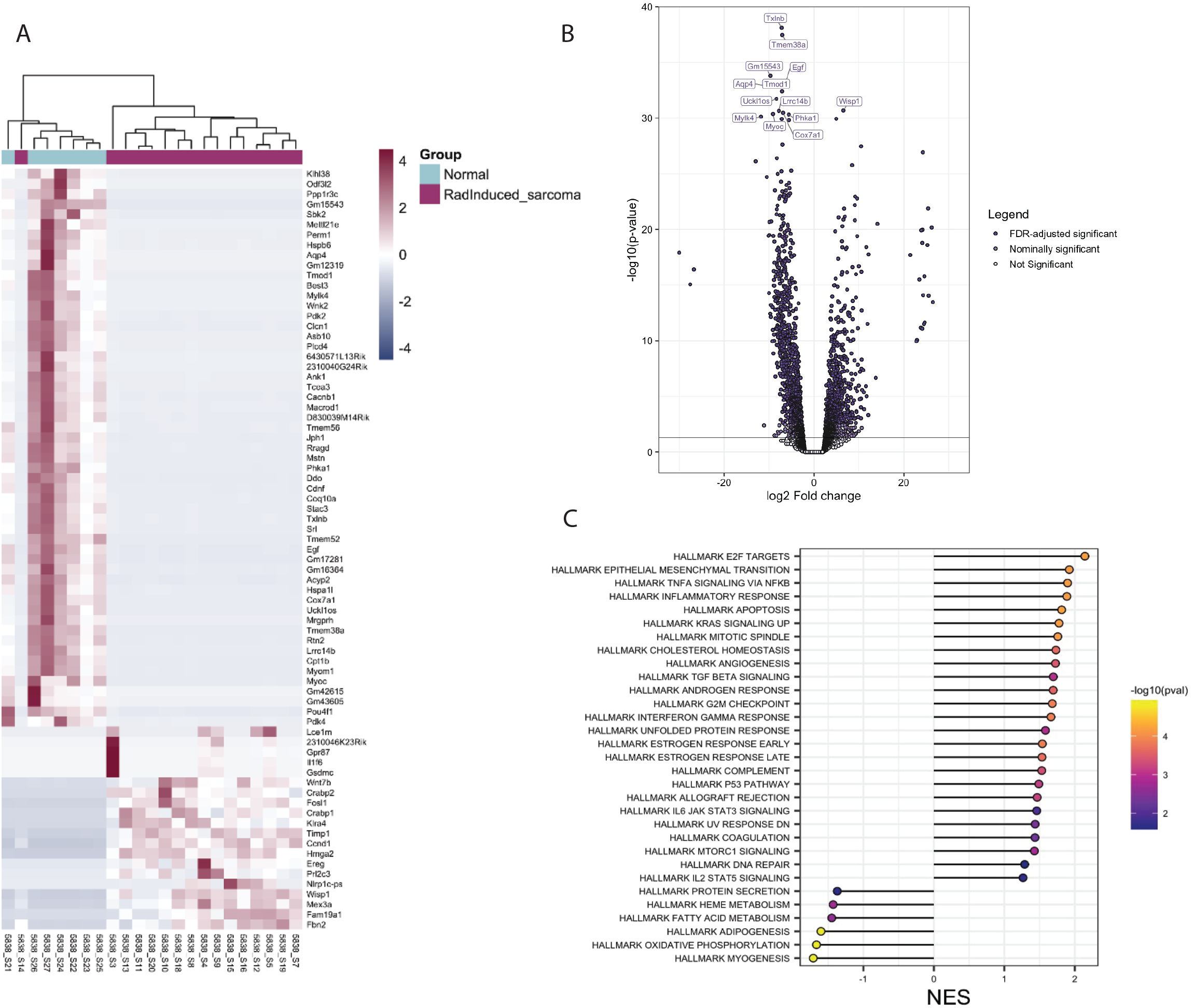
Radiation-induced sarcomas exhibit an increase in proliferative gene programs and a decrease in myogenic differentiation programs. (A) Heatmap of the top 75 differentially expressed genes in radiation-induced sarcomas (n=16) vs normal muscle (n=7). Genes (rows) are colored by scaled, normalized expression values. Both rows and columns are clustered. (B) Volcano plot of log2 fold change for all genes with base mean greater than 50 in tumor (n=16) vs normal (n=7). Labeled genes have a nominal -log10 p-value greater than or equal to 30. (C) Normalized enrichment scores for all false discovery rate (FDR) significant pathways for gene set enrichment analysis of Hallmark pathways. Points are colored by -log10 (nominal p-value).

Our previous WES data from a subset of the radiation-induced mouse sarcomas (n=7) revealed high copy number variations (CNV) (14) consistent with radiation-induced genetic damage in human tumors (33, 52, 53). We compared the specific oncogene amplification events identified by WES in each sarcoma with the gene expression data from the same tumor (Figure 5A-C and S8A). *Yap1, Met,* and *Cdk4* gene amplification resulted in significant transcriptional overexpression compared to normal muscle. In tumors where *Met* and *Cdk4* were not amplified (n=3) or the amplification status was unknown due to lack of WES (n=10) the expression of these genes was significantly upregulated, suggesting that activation of these oncogenes is selected for during tumor development either by amplification or alternative mechanisms (Figure 5B-C).

**Figure 5.**
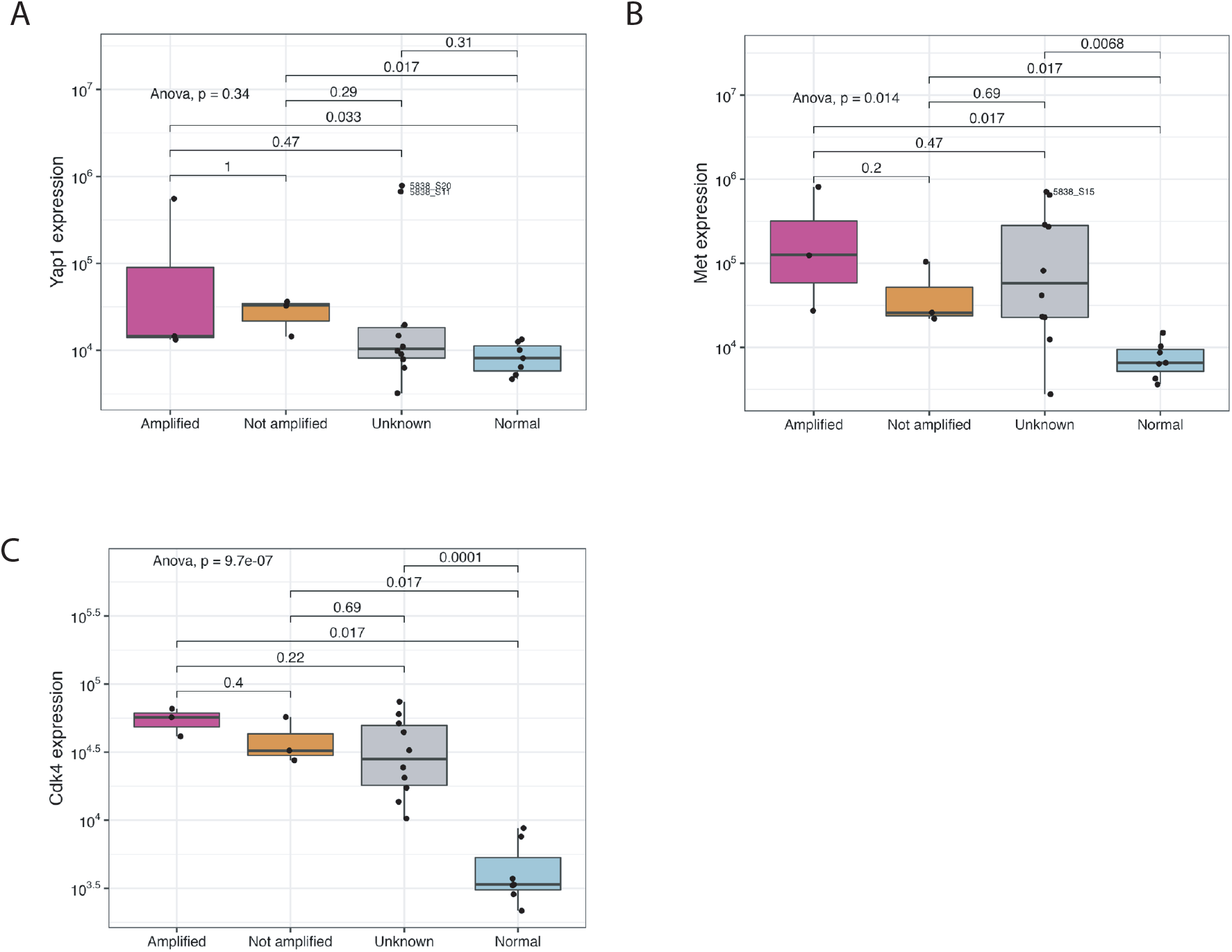
Comparison of copy number variation and expression of specific oncogenes. (A-C) Boxplot showing gene expression of *Yap1* (A), *Met* (B), and *Cdk4* (C) in radiation-induced sarcomas where the gene amplification status is known to be amplified (pink), not amplified (orange), or unknown (grey) compared to normal muscle (blue).

## Discussion

The strategy of reducing acute radiation-induced tissue injuries and adverse effects from other genotoxic therapies by blocking p53 promotes survival of damaged cells to preserve critical cellular functions in radiation-sensitive tissues (4). Therefore, inhibiting p53 during radiation therapy or radiation disaster scenarios has been proposed as an approach to prevent radiation injury. While blocking p53 during radiation exposure can protect the bone marrow from injury (13), our results in mice show that by temporarily blocking p53 during a single high-dose radiation exposure to a limb, late radiation effects, such as chronic tissue injury and radiation-induced sarcomagenesis can be exacerbated. Temporary knockdown of p53 during radiation exposure increased the development of sarcomas harboring genetic hallmarks of radiation damage. One possible explanation for these findings is that cells with severe DNA damage that would have died by p53-mediated cell death instead survived radiation injury due to p53 knockdown to become a sarcoma. As most of the mouse radiation-induced sarcomas mimicked UPS, which can arise from muscle satellite cells (12, 36), we used mice with a GFP reporter allele expressed from the endogenous Pax7 promoter to assess the impact p53 knockdown on the survival of muscle satellite cells after irradiation. We found that a temporary knockdown during a single fraction of 30 Gy to the limb preserved GFP+ cells in the limb, consistent with a model where inhibiting p53 preserves muscle satellite cells with DNA damage that have the potential to become a sarcoma. However, it is possible that temporarily blocking p53 during irradiation may increase radiation-induced sarcomas by other mechanisms.

Tissue sensitivity for radiation-induced tumorigenesis varies by cell type, radiation dose, fractionation (number of radiation exposures), volume of tissue irradiated, and other host factors such as age and germline mutations. The molecular mechanisms that govern the cellular responses to DNA damage, cell death, and cellular dynamics of tissue regeneration and remodeling impact tissue susceptibility to tumorigenesis (13, 54). The temporary and reversible p53 knockdown system provides a unique model to examine how blocking p53-mediated cell death from irradiation regulates radiation-induced cancer while avoiding permanent p53 loss, which is a well-known driver of radiation-induced cancer (29, 30, 55, 56). Using the same p53KD mouse model, we previously found that radiation-induced lymphoma development in mice following fractionated low dose total-body irradiation (TBI) is reduced when p53 expression is temporarily abrogated during irradiation (13). Temporary loss of p53 prevented cell death and improved bone marrow cell survival, thus ameliorating acute hematological toxicity and, surprisingly, lymphoma development, thereby improving overall survival of mice. In the TBI mouse lymphoma model, we demonstrated that temporary p53 knockdown reduces radiation-induced lymphomagenesis by limiting bone marrow cell death thus increasing cell competition for the thymic niche, which prevents the outgrowth of tumor-initiating cells in a non-cell autonomous mechanism. Niche competition is also the mechanism by which bone marrow transplantation prevents radiation-induced thymic lymphoma after TBI (57). Conversely, in this study we showed that temporarily blocking p53 during high-dose radiation promoted radiation-induced solid tumors, such as sarcomas, in a cell autonomous mechanism. Although our results are limited to an experimental system in which blocking radiation-induced cell death occurred by reducing p53, it is conceivable that these findings extend to other approaches to prevent or mitigate cell death from radiation injury. In this scenario, any strategy that promotes the survival of irradiated cells that were destined to die could increase the pool of potential tumor-initiating cells for a radiation-induced cancer. In the setting of a life-threatening radiation disaster, mitigating acute injury to increase the chance of survival would be worth the increased risk of a radiation-induced cancer years later. However, in the context of radiation therapy to treat a cancer, patients may not want an intervention that limits acute radiation toxicity while increasing the risk of radiation-induced tumorigenesis. Regardless, our results underscore the importance of preclinical testing of radiation protectors and mitigators in experiments with high-dose radiation similar to our study so that patients and their physicians will have information on the risk of a radiation modulator potentially exacerbating the risk of developing a radiation malignancy.

In patients, late adverse effects of radiotherapy are frequently irreversible and injuries to irradiated tissue can undergo protracted remodeling and healing phases (58, 59). Irradiated muscle exhibits impaired regenerative capacity despite satellite cell activation and vascular endothelial injury contributes to the development of fibrosis (60). In our study, chronic wounds after high-dose irradiation were characterized by inflammation, tissue fibrosis, tissue atrophy, occluded vasculature, and in severe cases, loss of the limb. Following 30 Gy irradiation we observed a significant increase in the development of chronic wounds (scores 1-3) in the p53 KD group compared to controls. We previously demonstrated that endothelial cells deficient in p53 are sensitized to late effects of radiation (41, 61, 62) which may contribute to reduced healing and increased wound severity. Notably, significant increases in chronic wounds in the p53KD group were only observed in the 40 Gy animals with a score of 2, suggesting that the higher dose of radiation was sufficient to overcome most of the protection afforded to tissues expressing normal levels of p53 following 30 Gy.

In addition to radiation-induced DNA damage through a cell autonomous process, our data suggest that high-dose irradiation may also promote tumor development as a consequence of prolonged tissue injury. Unresolved inflammation creates a permissive microenvironment for malignant conversion, where tumor cells are bathed in pro-growth, pro-remodeling, and pro-angiogenic signals (46, 63). Furthermore, tissue injury has been shown to accelerate tumor formation via promotion of epigenetic remodeling to induce a chromatin state that regulates gene expression programs favoring neoplastic commitment (49). Our prior studies demonstrated that muscle injury promotes sarcomagenesis though activation of the HGF-Met signaling axis to stimulate satellite cells (43, 44). Interestingly, our RNAseq data show that *Met* is upregulated in the majority of the radiation-induced sarcomas compared to normal muscle (n = 14). Further, radiation-induced sarcomas exhibited an enrichment of inflammatory signaling pathway genes which may result from developing within chronically inflamed tissue.

Others have examined the transcriptomes of human radiation-associated tumors but found no specific gene expression clustering associated with radiation exposure (34, 53). However, DNA sequencing of the radiation-associated human cancers reveals distinct radiation damage signatures that are also observed in the murine radiation-induced sarcomas (14, 30, 33, 53, 64).

In sum, the present study demonstrates that temporarily inhibiting p53 during high-dose irradiation of the limb promotes late effects of radiation, including sarcomagenesis. Our findings support a model for radiation sarcomagenesis that results from a combination of radiation-induced DNA damage and a permissive microenvironment from normal tissue injury that promote tumor development. In addition, these results suggest that blocking p53 during radiation therapy for a cancer might increase the risk of developing a radiation-associated malignancy.

## Acknowledgements

This work was supported by grants W81XWH-17-PRCRP-IA from the Department of Defense, U19AI067798 from the National Institute of Allergy and Infectious Diseases, and R35CA197616 from the National Cancer Institute (NCI) to DGK. This work was supported by the Whitehead Scholar award and K99CA212198 Award from the NCI to CL. We thank the Radiation Countermeasures Center for Research Excellence biostatistics core for their contributions to this work.

## Author Contribution Statement

AD, CL, YM, DK study conception and design. AD, CS, NW, JH, LL, YM, LC collected the data. ZL performed the statistical analysis of the mouse injury data. SS, JM, SL performed the analysis of the RNAseq data. DC performed the pathological analysis. AD, CL, CS, OL, YM, DK performed analysis and interpretation of data. AD, DK drafted manuscript. All authors reviewed and approved the manuscript.

---

**Figure S1.**
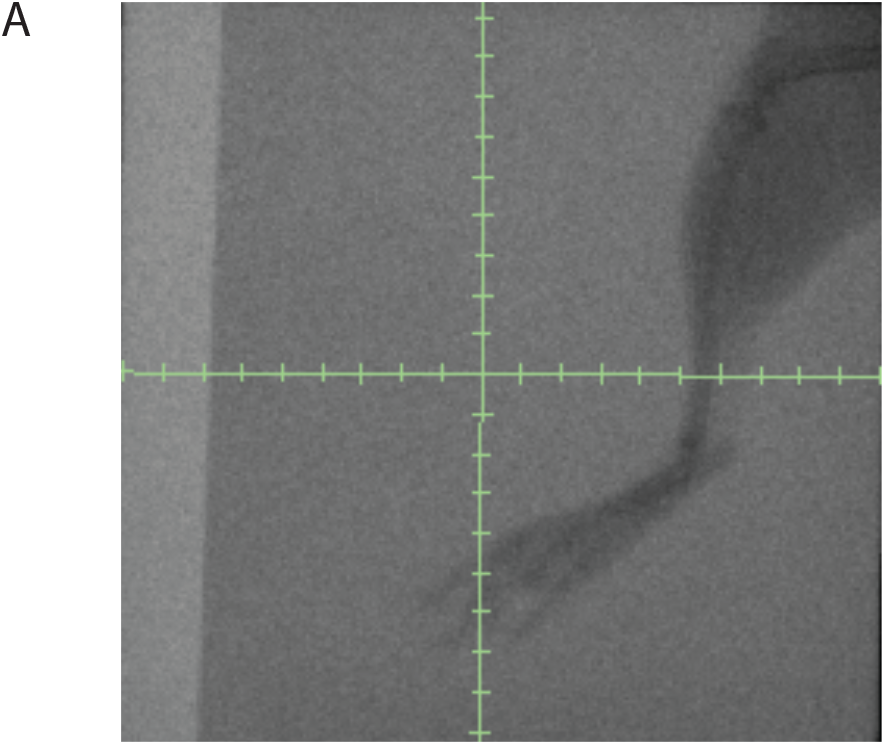
Mouse hind limb irradiation field. (A) Irradiations were performed using a small-animal image-guided irradiator and the target was defined using fluoroscopy with 40-kVp, 2.5-mA x-rays using a 2-mm aluminum filter. Representative image of the irradiation field is shown, which includes the whole hind limb.

**Figure S2.**
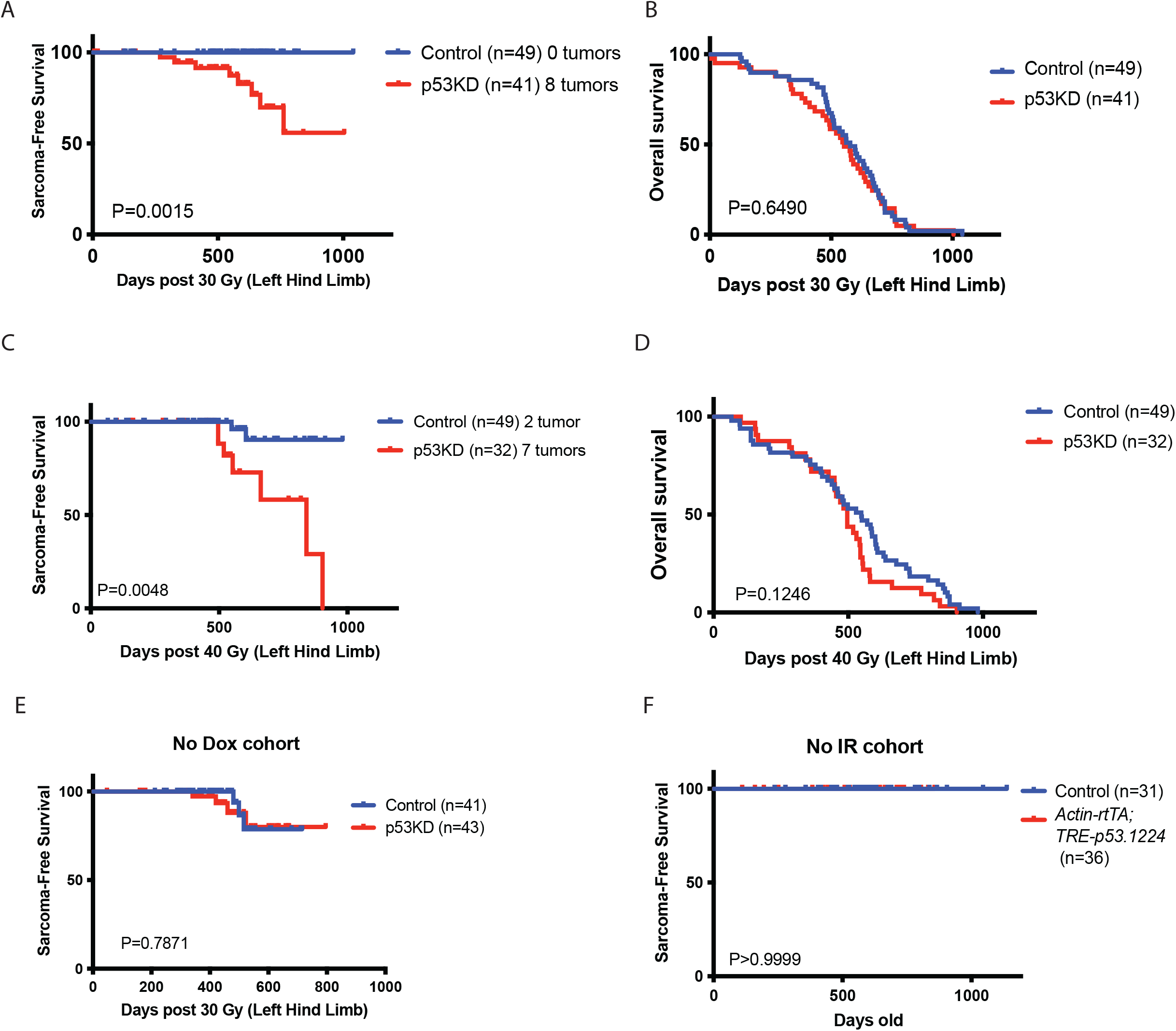
Temporary reduction in p53 expression during irradiation increases sarcomagenesis. (A) Kaplan-Meier curves show radiation-induced sarcoma-free survival of control and p53KD mice irradiated with 30 Gy to the hind limb. (B) Kaplan-Meier curves show overall survival of control and p53KD mice irradiated with 30 Gy to the hind limb. (C) Kaplan-Meier curves show radiation-induced sarcoma-free survival of control and p53KD mice irradiated with 40 Gy to the hind limb. (D) Kaplan-Meier curves show overall survival of control and p53KD mice irradiated with 40 Gy to the hind limb. (E) Kaplan-Meier curves show radiation-induced sarcoma-free survival of control and *Actin-rtTA; TRE-p53.1224* mice that did not receive dox and were irradiated with 30 Gy to the hind limb. (F) Kaplan-Meier curves show hind limb sarcoma-free survival of unirradiated control and *Actin-rtTA; TRE-p53.1224* mice. P – values are from log-rank tests.

**Figure S3.**
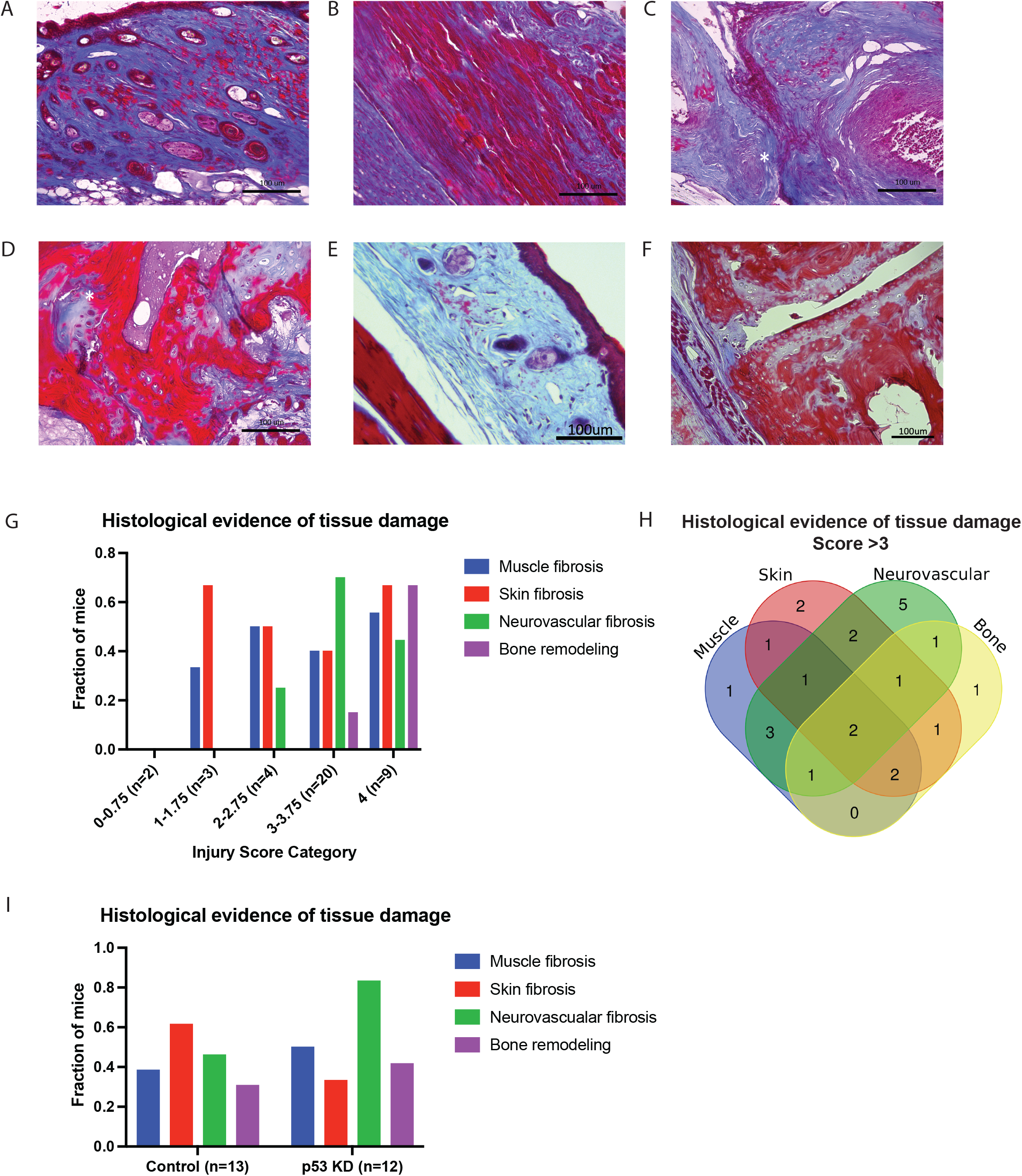
Histological examination of mouse radiation-induced injuries. (A-D) Representative histological image of trichrome-stained radiation-induced skin fibrosis (A), muscle fibrosis (B), neurovascular fibrosis (C, *fibrotic nerve), and bone remodeling (D, * bone-cartilage junction). (E-F) Representative images of uninjured muscle and skin (E) and bone and neurovascular structures (F). (G) Bar graph representing the fraction of mice from each injury score category with histological evidence of skin, muscle, neurovascular, and bone injuries. Note, some mice have injuries in more than one category. (H) Venn diagram representing the number of control and p53KD mice with injuries scores greater than 3 with injuries to multiple tissue types by histological analysis. (I) Bar graph representing the fraction of control and p53KD mice with injury scores greater that 3 with histological evidence of skin, muscle, neurovascular, and bone injuries. Note, some mice have injuries in more than one category.

**Figure S4.**
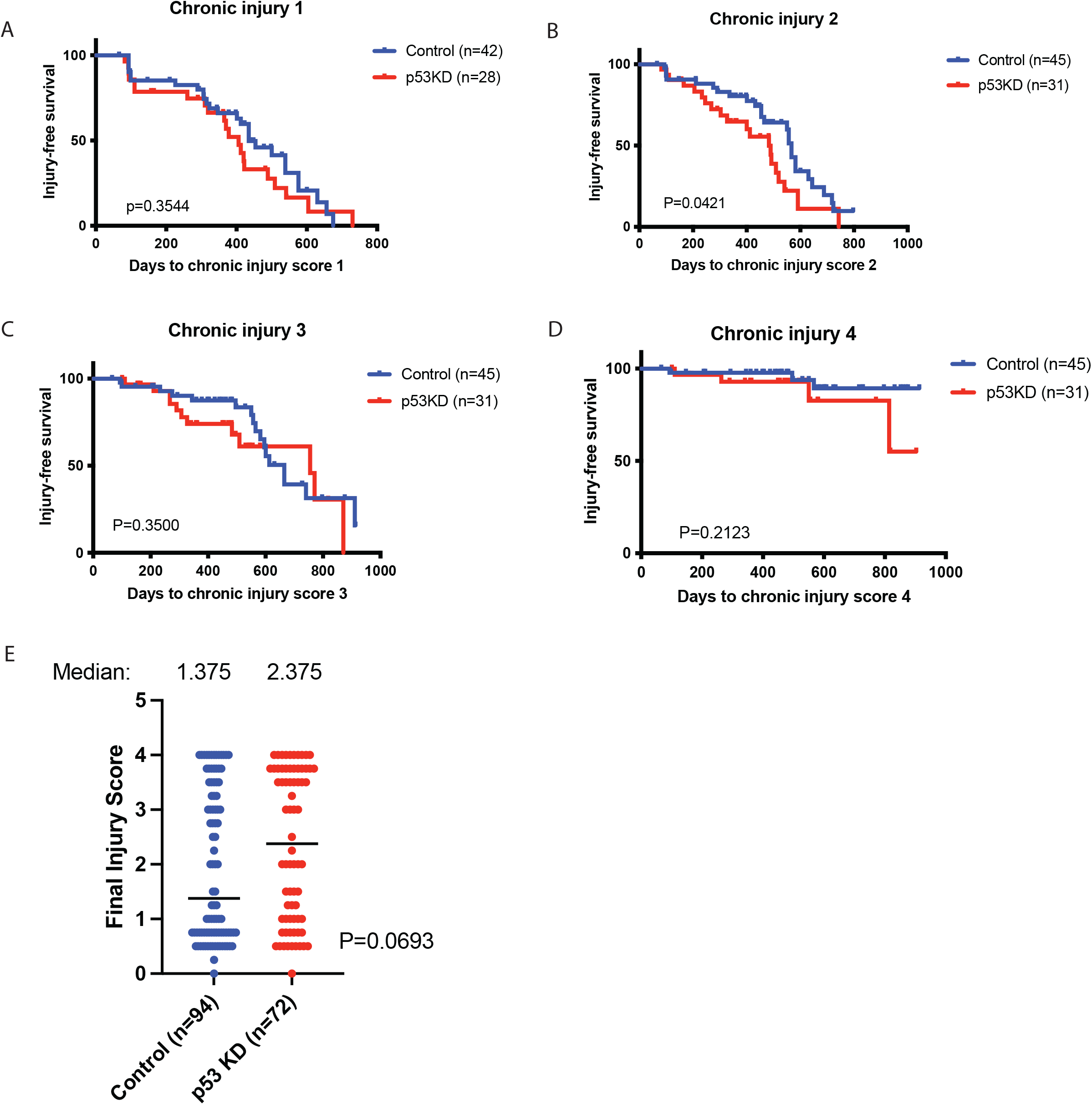
High dose irradiation induces chronic injuries in mouse hind limbs. (A-D) Kaplan-Meier curves show chronic injury-free survival from scores 1+ (A), 2+ (B), 3+ (C), or 4 (D) of control and p53KD mice irradiated with 40 Gy to the hind limb. P–value is from a log-rank test. (E) The final injury scores of the control and p53KD mice that received 30 or 40 Gy are plotted. P-value is from a T-Test.

**Figure S5.**
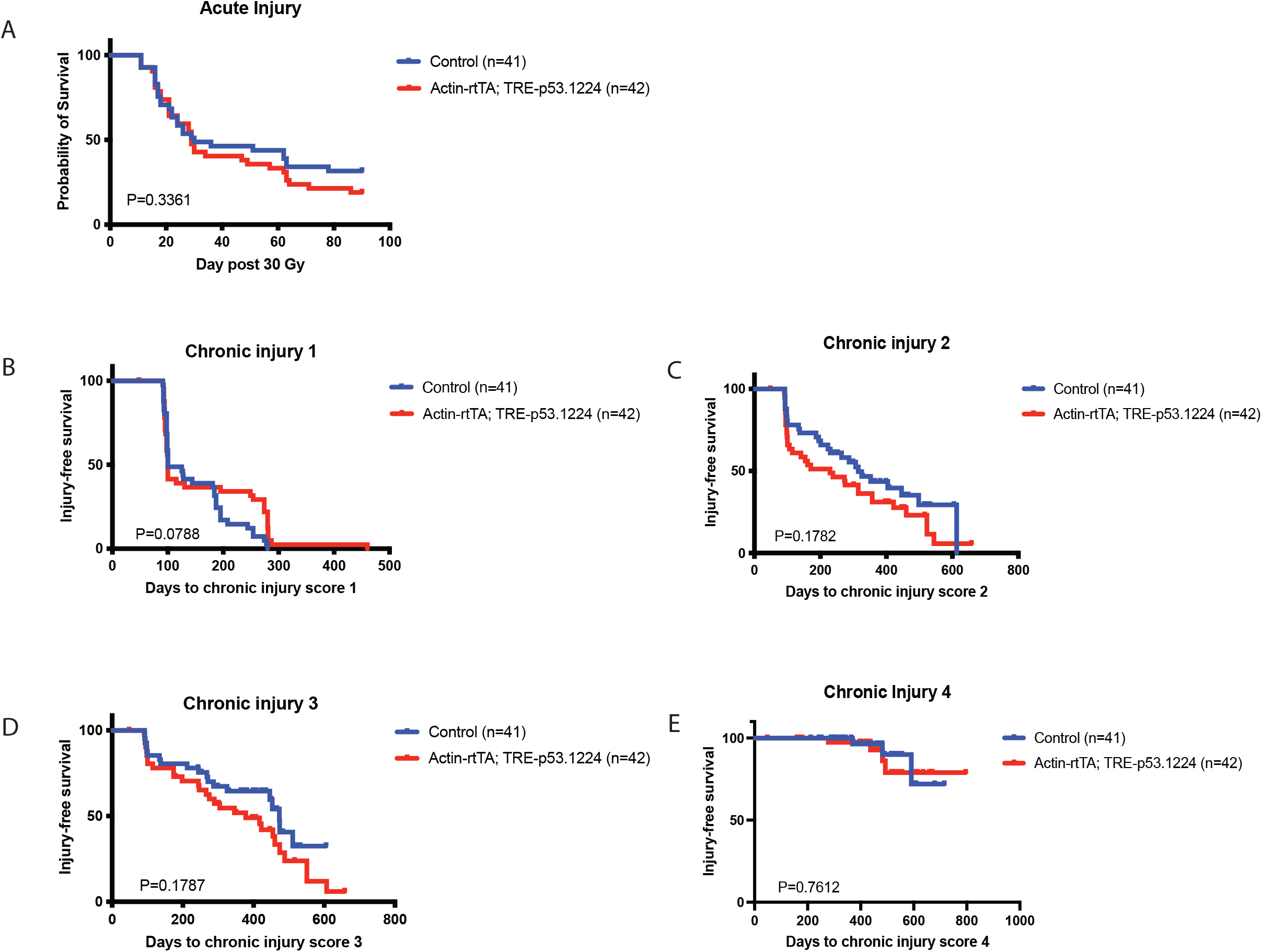
Mice without dox treatment sustained radiation-induced injuries in the hind limb. (A) Kaplan-Meier curves show acute injury-free survival (score 1+) of control and *Actin-rtTA; TRE-p53.1224* mice irradiated with 30 Gy to the hind limb without prior dox treatment. P-value is from a log-rank test. (B-E) Kaplan-Meier curves show chronic injury-free survival from scores 1+ (B), 2+ (C), 3+ (D), or 4 (E) of control and *Actin-rtTA; TRE-p53.1224* mice irradiated with 30 Gy to the hind limb without prior dox treatment. P-value is from a log-rank test.

**Figure S6.**
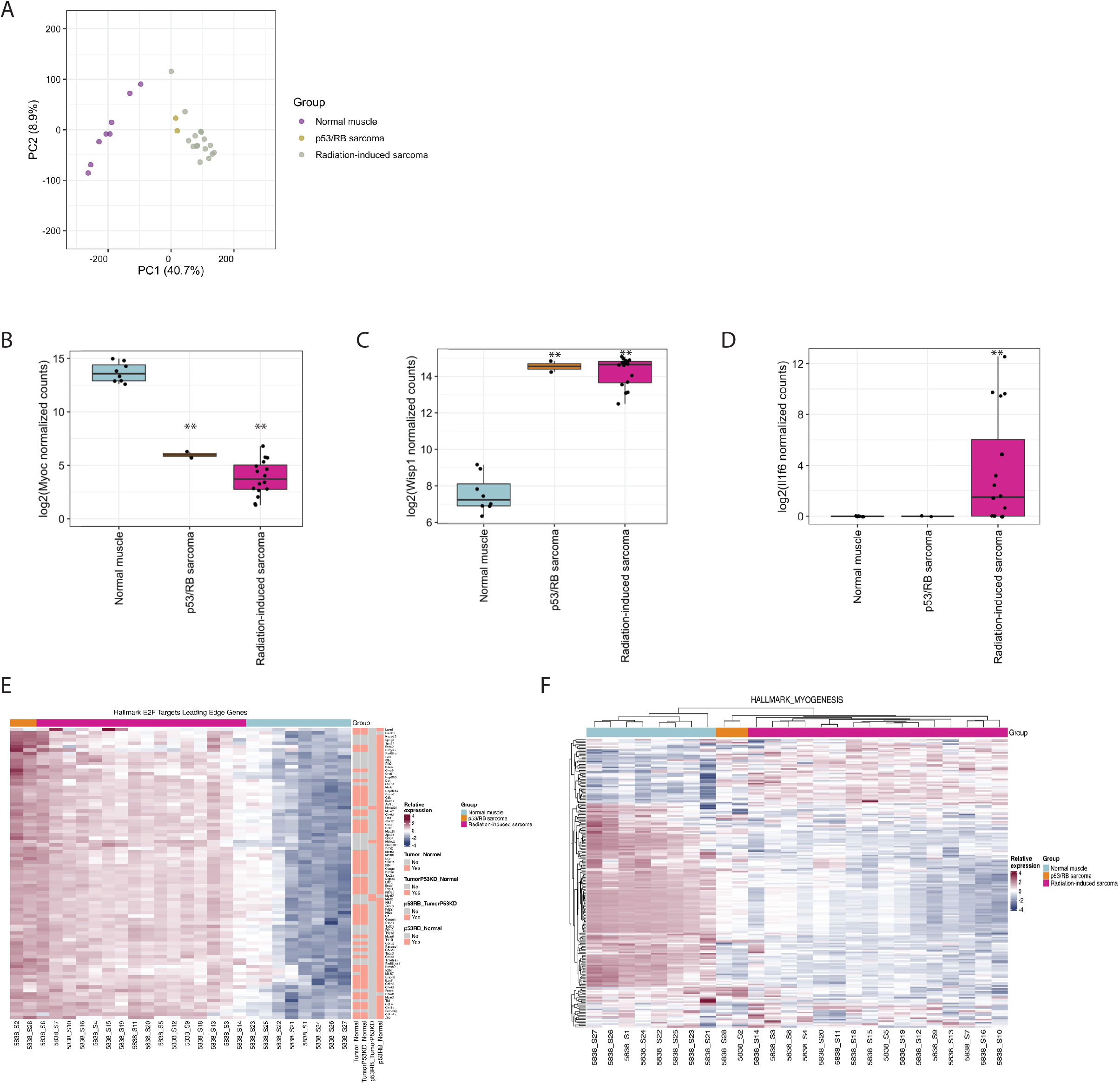
Gene expression analysis of radiation-induced sarcomas compared to normal muscles. (A) Principal component analysis of the mouse radiation-induced sarcomas compared to p53/RB sarcomas and normal muscles. (B) Boxplot of the differential expression of the *Myoc* gene in tumors compared to normal muscle. (C) Boxplot of the differential expression of the *Wisp1* gene in tumors compared to normal muscle. (D) Boxplot of the differential expression of the *Il1f6* gene in tumors compared to normal muscle. **P value >0.0001. (E) Heatmap of all genes in a leading-edge gene subset in the Hallmark E2F target pathway for any differential expression model. Expression is scaled relative to each row (gene), and samples and genes are hierarchically clustered. Genes in each row are annotated according to their membership in the leading edge of Hallmark E2F genes for each independent GSEA analysis (peach=in leading edge for a particular analysis, grey=not in leading edge). Samples are annotated by group (Normal, radiation-induced sarcoma, and p53/RB sarcoma samples). (F) Heatmap of all genes in the Hallmark Myogenesis target pathway. Expression is scaled relative to each row (gene) and genes are hierarchically clustered. Samples are annotated by group (Normal, radiation-induced sarcoma, and p53/RB sarcoma samples).

**Figure S7.**
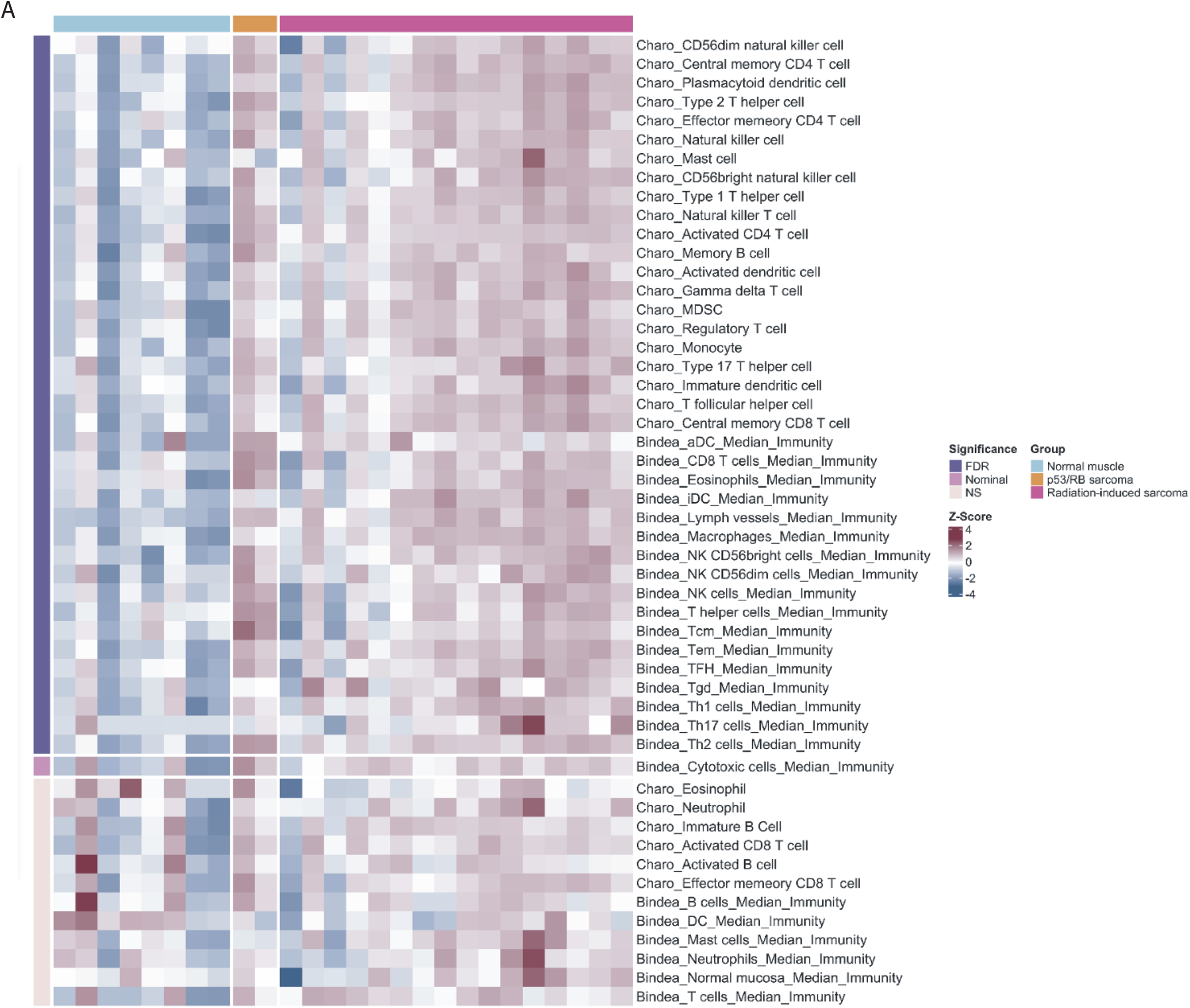
Immune infiltration analysis of radiation-induced sarcomas. (A) Heatmap showing radiation-induced sarcomas compared to normal muscles utilizing modules from Charoentong et al. (24) and Bindea et al. (23). Kruskal-Wallis test of scores was conducted across the groups. Module scores shown are Z-score transformed. Abbreviations: false discovery rate (FDR); not significant (NS)

**Figure S8.**
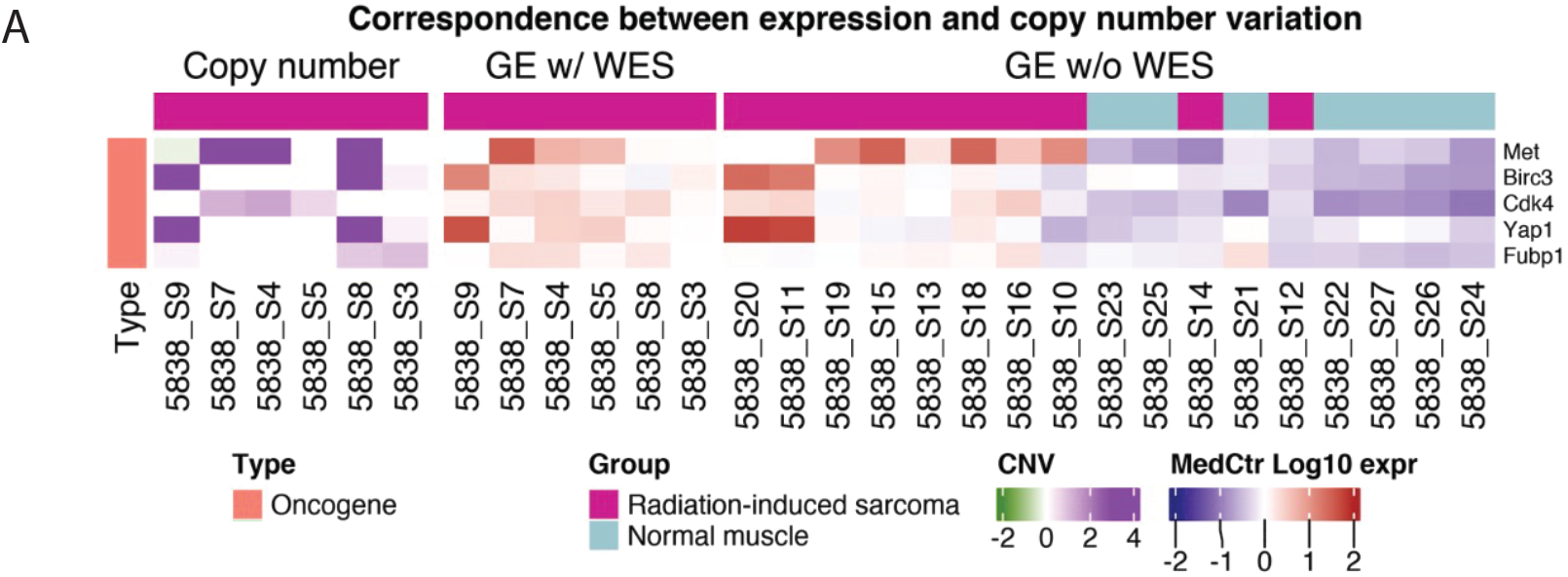
Correspondence between gene expression and CNV in radiation-induced sarcomas. (A) Heatmap of COSMIC oncogenes with CNV by WES (14) (left, purple/green scale) and the gene expression (GE) values (right, red/blue scale) from the same tumors (middle, GE w/ WES). The GE from the remaining radiation-induced tumors and normal muscles that were not subjected to WES are plotted (right).

**Table S1.**

|  | Mouse number | Tumor location | Genotype | Sex | Tumor latency | Injury score at death | LHL IR Dose | Treatment | Tumor type | RNAseq Sample ID |
| --- | --- | --- | --- | --- | --- | --- | --- | --- | --- | --- |
| 1 | 223030 | LHL | Control | M | 604 | 3.5 | 40 | Dox 10 days | UPS | 5838-S3 |
| 2 | 222933 | LHL | p53KD | M | 496 | 3.5 | 40 | Dox 10 days | UPS | 5838-S4 |
| 3 | 320114 | LHL | p53KD | F | 411 | 4 | 30 | Dox 10 days | UPS with myofibroblastic differentiation | 5838-S5 |
| 4 | 223181 | LHL | p53KD | M | 269 | 0.5 | 30 | Dox 10 days | UPS | 5838-S7 |
| 5 | 320006 | LHL | p53KD | M | 580 | 2.5 | 30 | Dox 10 days | UPS | 5838-S8 |
| 6 | 320239 | LHL | p53KD | M | 326 | 3.75 | 30 | Dox 10 days | UPS | 5838-S9 |
| 7 | 223178 | LHL | p53KD | M | 762 | 3.75 | 30 | Dox 10 days | UPS | 5838-S10 |
| 8 | 320359 | LHL | p53KD | M | 518 | 3.75 | 40 | Dox 10 days | UPS | 5838-S11 |
| 9 | 320258 | LHL | p53KD | F | 546 | 3.75 | 30 | Dox 10 days | UPS | 5838-S12 |
| 10 | 320177 | LHL | Control | F | 548 | 4 | 40 | Dox 10 days | UPS | 5838-S13 |
| 11 | 320085 | LHL | p53KD | M | 635 | 3.75 | 30 | Dox 10 days | Angiosarcoma | 5838-S14 |
| 12 | 320109 | LHL | p53KD | M | 669 | 4 | 30 | Dox 10 days | UPS | 5838-S15 |
| 13 | 320179 | LHL | p53KD | M | 662 | 4 | 40 | Dox 10 days | Giant cell UPS | 5838-S16 |
| 14 | 223032 | LHL | p53KD | M | 496 | 3.5 | 40 | Dox 10 days | UPS with myofibroblastic differentiation | NA |
| 15 | 320132 | LHL | p53KD | F | 903 | 4 | 40 | Dox 10 days | UPS | 5838-S18 |
| 16 | 320354 | LHL | p53KD | F | 840 | 4 | 40 | Dox 10 days | Leiomyosarcoma | 5838-S19 |
| 17 | 320358 | LHL | p53KD | M | 553 | 3.5 | 40 | Dox 10 days | UPS | 5838-S20 |

|  | Mouse number | Normal Muscle | Genotype | Sex | Days at death | RNAseq Sample ID |
| --- | --- | --- | --- | --- | --- | --- |
| 1 | 321103 | hind limb muscle | Actin-rtTA | M | 343 | 5838-S21 |
| 2 | 320679 | hind limb muscle | TRE-p53.1224 | M | 760 | 5838-S22 |
| 3 | 320703 | hind limb muscle | CMV-rtTA | M | 583 | 5838-S23 |
| 4 | 321281 | hind limb muscle | Actin-rtTA;TRE-p53.1224 | F | 243 | 5838-S24 |
| 5 | 320354 | hind limb muscle | Actin-rtTA;TRE-p53.1224 | F | 840 | 5838-S25 |
| 6 | 323338 | hind limb muscle | Actin-rtTA;TRE-p53.1224 | F | 117 | 5838-S26 |
| 7 | 323340 | hind limb muscle | Actin-rtTA | F | 117 | 5838-S27 |

LHL=Left Hind Limb

**Table S2.**

| Score | Description |
| --- | --- |
| 0 | Normal |
| 0.25 | Fifty-fifty doubtful if different from normal |
| 0.5 | Slight hair loss and/or very slight reddening |
| 0.75 | Definite but slight reddening +/- hair loss |
| 1 | Severe reddening with distended blood vessels or slight swelling |
| 1.25 | Severe reddening with white scales and/or severe swelling; red "papery" skin when healing |
| 1.5 | Moist breakdown of one small area (usually on bottom of foot first) with scaly appearance |
| 1.75 | Moist desquamation in more than one small area or one slightly larger area (tips of toes stuck together with no other breakdown when healing) |
| 2 | Breakdown of larger area and/or toes stuck together; possibly moist in places |
| 2.25 | Breakdown in one-third area of foot |
| 2.5 | Breakdown of one-half area of foot (usually first on bottom) |
| 2.75 | Breakdown of about two-thirds area of foot |
| 3 | Breakdown of most of skin of foot, possible slight moist exudate; possible paralysis |
| 3.25 | Breakdown of entire skin of foot with slight moist exudate |
| 3.5 | Breakdown of entire skin with severe moist exudate; may be stuck to body fur |
| 4 | Loss of foot |

